# Co-occurrence of *kdr* mutations V1016I and F1534C in pyrethroid-resistant *Aedes aegypti* (Diptera: Culicidae) populations from Costa Rica

**DOI:** 10.1101/704767

**Authors:** Aryana Zardkoohi, David Castañeda, Carmen Castillo, Juan C Lol, Francisco Lopez, Rodrigo Marín Rodriguez, Norma Padilla

## Abstract

*Aedes aegypti* (Linnaeus, 1762) is considered the most important mosquito vector species for several arboviruses (e.g., dengue, chikungunya, Zika) in Costa Rica. The main strategy for the control and prevention of *Aedes*-borne diseases relies on insecticide-based vector control. However, the emergence of insecticide resistance in the mosquito populations present a big threat for the prevention actions. The characterization of the mechanisms driving the insecticide resistance in *Ae. aegypti* are vital for decision making in vector control programs. Therefore, we analyzed the voltage-gated sodium channel gene for the presence of the V1016I and F1534C *kdr* mutations in pyrethroid-resistant *Ae. aegypti* populations from Puntarenas and Limon provinces, Costa Rica. The CDC bottle bioassays showed that both Costa Rican *Ae. aegypti* populations were resistant to permethrin and deltamethrin. In the case of *kdr* genotyping, results revealed the co-occurrence of V1016I and F1534C mutations in permethrin and deltamethrin-resistant populations, as well as the fixation of the 1534C allele. Therefore, our findings make an urgent call to expand the knowledge about the insecticide resistance status and mechanisms in the Costa Rican populations of *Ae. aegypti* which must be a priority to develop an effective resistance management plan.

## Introduction

The region of the Americas has experienced the emergence and re-emergence of *Aedes*-borne diseases (ABD; e.g., dengue, chikungunya, Zika) (Weaver and Reisen 2010, Espinal et al. 2019), with the mosquito *Aedes aegypti* (Diptera: Culicidae, Linnaeus) widely distributed across the region (Leta et al. 2018). Costa Rica reported one of the highest numbers of confirmed cases of Zika virus (ZIKV) during the Zika emergency from 2015-2018 in the Central America region (PAHO 2018). Despite control efforts, ABD persists in Limon and Puntarenas provinces, where the highest number cases of ZIKV of Costa Rica were reported (Sanchez et al. 2019).

One of the main strategies for the control and prevention of ABD in Costa Rica relies on insecticide-based vector control, targeting *Aedes* populations. For the adult populations, several pyrethroids have been used sequentially (deltamethrin, cyfluthrin, cypermethrin) (Marín Rodriguez et al. 2009, Bisset et al. 2013), and currently permethrin and lambda-cyhalothrin are being used (Ministerio de Salud de Costa Rica 2019). However, focal studies have reported a decrease in the susceptibility against pyrethroids in *Ae. aegypti* from Puntarenas and Limon (Bisset et al. 2013, Calderón-Arguedas and Troyo 2014, 2016). Moreover, biochemical and synergists assays suggest that carboxylesterases and cytochrome P450s could be involved in pyrethroid resistance. Besides from biochemical activity, one of the main mechanisms of pyrethroid resistance in insects is the target-site insensitivity (Moyes et al. 2017), which has been poorly described in Costa Rica (Chaves et al. 2015). Non-synonymous mutations on the knockdown resistance (*kdr*) region of the voltage-gated sodium channel (*VGSC*) gene are responsible for this pyrethroid insensitivity (Du et al. 2016). A study published in 2003 reported seven novel mutations in *Ae. aegypti* strains from Asia, Africa, the Caribbean, South America and Oceania (Brengues et al. 2003). Although many non-synonymous mutations have been discovered in the *Ae. aegypti* VGSC gene, only five have been associated with pyrethroid resistance: V410L, S989P, I1011M, V1016G, and F1534C (Du et al. 2013, Hirata et al. 2014, Haddi et al. 2017).

Molecular screening of the Latin American populations of *Ae. aegypti* evidenced the emergence of the *kdr* mutation V1016I in the region (Saavedra-Rodriguez et al. 2007) and its co-occurrence with the I1011M mutation, which was recently found in Panama (Murcia et al. 2019). The first report of the F1534C mutation was detected in 2011 in the *Ae. aegypti* population from Thailand (Yanola et al. 2011). Since then, the F1534C mutation has been documented worldwide (Linss et al. 2014, Kushwah et al. 2015, Hamid et al. 2018, Maestre-Serrano et al. 2019, Sombié et al. 2019), except from Central America. To the best of our knowledge, *kdr* mutations have not been studied in Costa Rican areas where pyrethroid resistance has been reported for *Ae. aegypti*. In order to fill the gap about the mechanisms associated with insecticide resistance, we analyzed the *VGSC* gene for the presence of *kdr* mutations in pyrethroid-resistant *Ae. aegypti* populations from Limon and Puntarenas.

## Material and Methods

### Mosquito sampling and rearing

*Aedes* mosquito eggs were collected as part of the National Entomological Surveillance of the Ministry of Health from Costa Rica in two provinces (Figure 1): Puntarenas (in the Pacific coast) and Limon (in the Caribbean coast). For each site, 50 ovitraps were placed and visited weekly to collect the mosquito eggs during May 2017. Eggs collected from the ovitraps for each site were pooled, hatched and the larvae reared into adults under insectary conditions (28 ± 2°C; 70 ± 10% RH, and 12:12 photoperiod). A F1 generation was obtained to get enough individuals to perform the bioassays. All emerged adults were provided access to cotton pads soaked in 7% sugar solution until bioassays.

**Fig 1.**
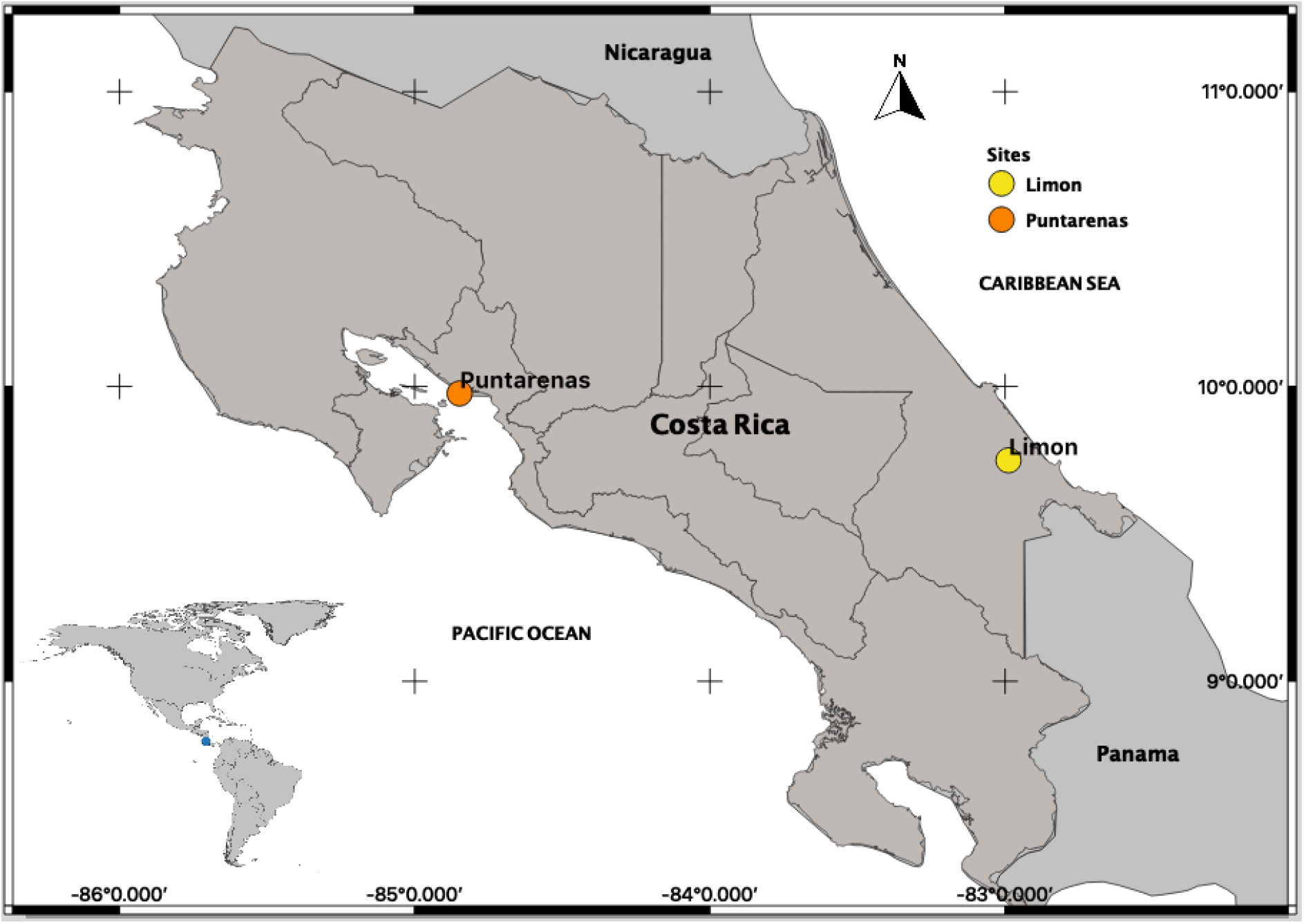
*Aedes* mosquito collection sites. A map of Costa Rica shows the location of Puntarenas (in the Pacific coast) and Limon (in the Caribbean coast) provinces.

### Adult bioassays

Insecticide susceptibility against permethrin (type I pyrethroid) and deltamethrin (type II pyrethroid) in the Costa Rican *Ae. aegypti* populations were assessed with the CDC bottle bioassay (Brogdon and McAllister 1998). From each population, a total of 100 F1 females (2-5-days old, non-blood-fed) were exposed to diagnostic dose of deltamethrin (10 μg/bottle) or permethrin (15 μg/bottle) for 30 min as diagnostic time. The New Orleans strain was used as insecticide-susceptible reference strain to support the appropriate performance of the bioassays. Based on the incapacitation rate at 30 min, each population was classified according to the standard WHO criteria as follows (WHO 2016): a population with an incapacitation rate >98% was classified as susceptible (S), an incapacitation rate between 90-98% suggested possible resistance (PR) requiring further tests, and an incapacitation rate <90% indicated confirmed resistance (R). Resistant and incapacitated mosquitoes were preserved individually at −20°C for molecular assays.

### *Kdr* genotyping

Genomic DNA (gDNA) was extracted from permethrin and deltamethrin-resistant mosquitoes using DNAzol (Thermo Fisher Scientific) according to manufacturer’s instructions with modifications. Briefly, 100 uL of DNAzol was used to homogenize a whole mosquito and DNA pellets were resuspended in 50 uL of DEPC water. Subsequently, *kdr* genotyping was performed to detect the presence of V1016I and F1534C mutations in the pyrethroid-resistant mosquitoes. The genotype of each mutation was determined by a melt curve analysis of a real-time PCR using allele-specific primers for the V1016I mutation (Val1016f, Ile1016f and Ile1016r) reported by Saavedra-Rodriguez et al. (2007), and for the F1534C mutation (F1534-f, C1534-f and CP-r) reported by Yanola et al. (2011).

PCR reaction for the V1016I mutation contained 10 μL of 2X PowerUp SYBR Green Master Mix (Thermo Fisher Scientific), 0.5 μL of each primer (50 pmol/μL), 1 μL of gDNA (5 to 15 ng) in a final volume of 20 μL. The PCR conditions were as followed: 95°C for 2 min, 40 cycles of 95°C for 10 sec, 60°C for 10 sec, and 72°C for 30 sec, with a final stage of 10 sec at 95°C. The melt curve conditions were 65°C to 95°C with an increase of 0.2°C each 10 sec. Based on the melting temperatures, samples were considered as mutant homozygous if they presented a unique peak of 79°C, wildtype homozygous if they presented a unique peak of 85°C, and heterozygous if they presented both of the previous peaks. In the case of F1534C mutation, each reaction contained 10 μL of 2X PowerUp SYBR Green Master Mix (Thermo Fisher Scientific), 0.066 μL of C1534-f primer (50 pmol/μL), 0.2 μL of primers F1534-f (50 pmol/μL) and CP-r (50 pmol/μL), 1 μL of gDNA (5 to 15 ng) in a final volume of 20 μL. The PCR conditions were as followed: 95°C for 3 min, 40 cycles of 95°C for 10 sec, 57°C for 10 sec, and 72°C for 30 sec, with a final stage of 10 sec at 95°C. The melt curve conditions were 65°C to 95°C with an increase of 0.5°C every 5 sec. Based on the melting temperatures, samples were considered as mutant homozygous if they presented a unique peak of 85°C, wildtype homozygous if they presented a unique peak of 80°C, and heterozygous if they presented both of the previous peaks. Previously genotyped *Ae. aegypti* by sequencing were used as positive controls for the PCR assays. The genotypic and allelic frequencies were calculated for each insecticide and population.

### Sequencing of the *VGSC* gene

To validate the genotypes obtained from the *kdr* genotyping, two different PCR assays were used to amplify a partial fragment of domain IIS6 and domain IIIS6 of the *VGSC* gene in a subsample of 13 mosquitoes which represents the different genotypes identified by the real-time PCR. The first assay amplified the domain IIS6 using the primers IIS5-6 F and IIS5-6 R (spaning the codons 989, 1011 and 1016) originally reported by Al Nazawi et al. (2017). Each PCR reaction contained 1X Colorless GoTaq Flexi Buffer, 1.5 mM MgCl2, 0.2 mM dNTPs, 2.5 μM of each primer (IIS5-6F and IIS5-6R), 1.25 U of GoTaq Hot Start Polymerase (Promega), and 1.25 ng of gDNA in a total volume of 50 μL. The cycling parameters included an initial step of denaturation at 95°C for 2 min, followed by 40 cycles of 94°C for 30 sec, 50°C for 30 sec, and 72°C for 1 min, with a final extension step of 10 min at 72°C. For the second assay, a region of the domain IIIS6 was amplified using the primers AaNa31F and AaNa31R (spaning the codon 1534) reported by Harris et al. (2010). Each PCR reaction contained 1X Colorless GoTaq Flexi Buffer, 1.5 mM MgCl2, 0.2 mM dNTPs, 0.5 μM of each primer (AaNa31F and AaNa31R), 2.5 U of GoTaq Hot Start Polymerase (Promega), and 5 ng of gDNA in a total volume of 50 μL. The cycling parameters included an initial denaturation at 95°C for 2 min, followed by 35 cycles of 94°C for 30 sec, 59°C for 30 sec, and 72°C for 1 min, with a final extension step of 10 min at 72°C. Both PCR assays were performed using the SimpliAmp Thermal Cycler (Applied Biosystems).

All PCR products were visualized on a 2% agarose gel stained with ethidium bromide under UV light. Then, 50 uL of PCR product of each sample was purified with SpinPrep PCR Clean-Up Kit (EMD Millipore Corp) according to manufacturer’s instructions. The PCR products purified were sequenced by Macrogen Inc. (Maryland, USA) with the primers used in the PCR assays. Sequences were assembled and aligned using MEGA 7 (Kumar et al. 2016), and all codons were analyzed for non-synonymous substitutions.

## Results and Discussion

The results of the CDC bottle bioassays are presented in Table 1. Incapacitation rates varied between 35 and 87% for deltamethrin and permethrin, confirming resistance to these pyrethroids in both Costa Rican *Ae. aegypti* populations. Puntarenas exhibited lower incapacitation rate (between 35 and 62%) for both insecticides compared with Limon (between 79 and 87%). Incapacitation rates for permethrin (between 35 and 79%) were lower than deltamethrin (between 62 and 87%) in the two *Ae. aegypti* populations from Costa Rica. A previous study in 2013, found suspected resistance to deltamethrin (incapacitation rate of 90.5%) in a Puntarenas *Ae. aegypti* population (Bisset et al. 2013). Therefore, our results suggest that deltamethrin resistance persists as well as an increase in the deltamethrin resistance from Puntarenas *Ae. aegypti* population.

**Table 1.** Pyrethroid susceptibility in Costa Rican *Aedes aegypti* populations assessed by CDC bottle bioassay.

| Population | n | % incapacitation [95% CI] | Status |
| --- | --- | --- | --- |
| Permethrin |  |  |  |
| Puntarenas | 100 | 35 [16.23-53.77] | R |
| Limon | 100 | 79 [77.04-80.96] | R |
| Deltamethrin |  |  |  |
| Puntarenas | 100 | 62 [54.49-69.51] | R |
| Limon | 100 | 87 [85.04-88.96] | R |

As part of the insecticide resistance management, early detection of insecticide resistance mechanisms is critical for decision making in vector control programs (Corbel et al. 2019). To our knowledge, this is the first study to found and report a co-occurrence of both F1534C and V1016I mutations in resistant populations from Costa Rica (Table 2). The three different possible genotypes (V/V, V/I and I/I) for the *kdr* mutation V1016I were found in Puntarenas and Limon. However, for the 1534 codon, the homozygous mutant genotype C/C was the only one detected in the 72 mosquitoes analyzed. This result suggests that the mutation could be fixed in the field populations. The *kdr* allele frequency for 1016I varied between 0.167 and 0.980, and the *Ae. aegypti* population from Puntarenas showed a higher allele frequency than Limon. In addition, DNA sequencing confirmed the presence of all genotypes observed by the allele-specific real-time PCR in the two Costa Rican *Ae. aegypti* populations.

**Table 2.** Genotypic and allelic frequencies of the V1016I and F1534C *kdr* mutations in pyrethroid-resistant mosquitoes from Costa Rican *Aedes aegypti*.

| Population | n | V1016I genotypes |  |  | F1534C genotypes |  |  | *kdr* allele frequency |  |
| --- | --- | --- | --- | --- | --- | --- | --- | --- | --- |
|  |  | V/V | V/I | I/I | F/F | F/C | C/C | 1016I | 1534C |
| Permethrin-resistant |  |  |  |  |  |  |  |  |  |
| Puntarenas | 13 | 1 | 5 | 7 | 0 | 0 | 13 | 0.731 | 1 |
| Limon | 21 | 16 | 3 | 2 | 0 | 0 | 21 | 0.167 | 1 |
| Deltamethrin-resistant |  |  |  |  |  |  |  |  |  |
| Puntarenas | 25 | 0 | 1 | 24 | 0 | 0 | 25 | 0.980 | 1 |
| Limon | 13 | 7 | 3 | 3 | 0 | 0 | 13 | 0.347 | 1 |

The *kdr* mutation V1016I was originally reported as a widespread mutation across many Latin American *Ae. aegypti* populations, including a population from Guanacaste province, Costa Rica (Saavedra-Rodriguez et al. 2007). Although, functional conformation experiments do not confirm the role of the V1016I mutation in pyrethroid resistance, many studies have correlated the presence of this mutation to phenotypic resistance (Deming et al. 2016, Brito et al. 2018, Maestre-Serrano et al. 2019, Ryan et al. 2019). In contrast, *kdr* mutation F1534C has been strongly associated with DDT and pyrethroid resistance and confirmed by functional experiments in *Xenopus oocytes* (Du et al. 2013). Higher levels of pyrethroid resistance in mosquitoes have been observed in populations with the co-occurrence of the mutations V1016I and F1534C in other countries (Aponte et al. 2013, Kawada et al. 2014, Chapadense et al. 2015, Dusfour et al. 2015, Sombié et al. 2019). This co-occurrence has been observed approximately in the 97% of mosquitoes analyzed from Puntarenas, whereas a 32% of the mosquitoes from the province of Limon presented both mutations V1016I and F1534C (Table 2). These results could explain the different levels of pyrethroid resistance between the two *Ae. aegypti* populations analyzed (Table 1). In agreement with Linss et al. (2014), the V1016I mutation was not present without the F1534C mutation. Based on recent evidence, fixation of F1534C mutation in the populations analyzed represents a major threat for the insecticide-based vector control in Costa Rica (Vera-Maloof et al. 2015). Since this fixated mutation could favor the emergence of other *kdr* mutations, like the V1016I. Multiple *kdr* mutations could cause an additive effect increasing the levels of insecticide resistance and decreasing significantly the efficiency of vector control programs (Li et al. 2015, Plernsub et al. 2016, Saavedra-Rodriguez et al. 2018).

Overall, these results evidenced the presence of *kdr* mutations in permethrin and deltamethrin-resistant *Ae. aegypti* individuals from Costa Rica as well as the first report of the F1534C mutation and its co-occurrence with V1016I. Therefore, our findings make an urgent call to expand the knowledge about the insecticide resistance status and mechanisms in the Costa Rican populations of *Ae. aegypti* which must be a priority to develop an effective resistance management plan.

## Acknowledgments

We thank Dr. Lissette Navas, INCIENSA’s President, for facilitating the development of this research. This work was supported by Cooperative Agreement Number U51GH000970 funded by the Centers for Disease Control and Prevention (CDC). Its contents are solely the responsibility of the authors and do not necessarily represent the official views of the Centers for Disease Control and Prevention from USA or the Department of Health and Human Services from the Costa Rica’s Ministry of Health. The authors declare that they have not conflicts of interest.

